# Mediodorsal thalamus of alcohol-dependent mice shows genetic and physiological adaptations and alcohol-biased calcium signaling

**DOI:** 10.1101/2024.10.28.620696

**Authors:** Jayden C. Martin, Kaitlin C. Reeves, Kathryn A. Carter, Matthew Davis, Amelia Schneider, Elizabeth Meade, Christina L. Lebonville, Sudarat Nimitvilai-Roberts, Michaela Hoffman, John J. Woodward, Gabriel Loewinger, Rachel J. Smith, Marcelo F. Lopez, Howard C. Becker, Patrick J. Mulholland, Jennifer A. Rinker

## Abstract

Alcohol Use Disorder (AUD) is a significant health concern characterized by cognitive dysfunction and an inability to control alcohol intake leading to severe social and health consequences. It is crucial to uncover neuroadaptations and cellular mechanisms responsible for poor decisions surrounding alcohol drinking. Although the mediodorsal thalamus (MD) is an essential brain region for cognitive function and reward-guided choices, the effects of alcohol dependence on MD neuroadaptations and how dependence alters MD activity during choice behaviors for alcohol over natural rewards (i.e., sucrose) are not well understood. Genetic and physiological adaptations in the MD were assessed in mice treated with the chronic intermittent ethanol (CIE) exposure model of dependence, which increased alcohol intake and preference during choice sessions for water or sucrose. Results indicated that CIE exposure induced changes in transcript expression and excitability of MD neurons. Enrichment analysis of alcohol-sensitive genes revealed dysregulation of genes that control glial function and axonal myelination. Fiber photometry was used to record MD calcium signals in CIE mice during choice drinking sessions for alcohol and either water or sucrose. Results demonstrated that MD activity was elevated at the start of and after licking bouts for alcohol, water, or sucrose, although the signal for alcohol was significantly higher than signal for other solutions and this effect persisted after induction of dependence. These findings demonstrate that CIE exposure causes genetic and physiological neuroadaptations in the MD that coincide with alcohol-biased behaviors, with the MD uniquely responsive to alcohol over other solutions.

## 1. Introduction

Alcohol Use Disorder (AUD) is characterized by excessive alcohol consumption and the inability to control drinking behavior. AUD is the most common substance use disorder globally and affects approximately 8% of the population (Ariesen et al., 2023). Since 1990, binge drinking prevalence has sharply increased, while lifetime abstinence rates have declined (Manthey et al., 2019). The rates of excessive alcohol consumption are predicted to rise over the next decade, further intensifying alcohol-related health issues and diminishing quality of life. Individuals with AUD frequently display poor decision-making and impaired behavioral flexibility, often manifesting as a choice bias for alcohol over other responsibilities and natural rewards, e.g., work, family, alternative non-drug rewards. Deficits in behavioral flexibility—the ability to adjust actions based on changes in reward value—is a critical characteristic of alcohol and substance use disorders (Wicker et al., 2018). Impaired behavioral flexibility is often associated with mood disorders, anxiety, and their psychiatric comorbidities, like AUD. Individuals with AUD show increased alcohol-approach associations on an implicit association task, and this alcohol-biased behavior is associated with increased lifetime alcohol use and fewer days of abstinence (Wiers et al., 2017). This biased choice for alcohol has also been modeled in alcohol dependent and non-dependent rodents, whereby rodents choose alcohol over social interaction and sweetened solutions (Augier et al., 2018; Augier et al., 2023; Lopez & Becker, 2025; Marchant et al., 2023; Russo et al., 2018). Identifying regions and mechanisms responsible for the shift toward alcohol biased behaviors is, therefore, essential for developing therapeutic interventions for AUD.

The mediodorsal thalamus (MD) is a crucial region of interest in AUD research due to its role in memory, cognition, and reward-guided behavior (Parnaudeau et al., 2018; Savage et al., 2021). The dense interconnected circuitry between the MD and prefrontal cortex (PFC) influences higher-order cognitive functions, such as reward processing and decision-making, including choice for high vs. low rewards (Ding et al., 2024). MD lesions impaired the updating of reward value, as seen in reinforcer devaluation tasks that assess how changes in the value of a reward influence choice behavior (Wicker et al., 2018). In studies with non-human primates, MD lesions resulted in cognitive deficits comparable to those following PFC damage, especially in tasks involving flexible reward updating and reinforcement devaluation (Alcaraz et al., 2018; Chakraborty et al., 2016). Additionally, the MD functions as both a simple relay structure and a dynamic integrator, stabilizing task-relevant PFC activity and facilitating top-down signaling (Bolkan et al., 2017). Thalamocortical loops involving the MD support adaptive decision-making, behavioral flexibility, and cognitive domains that are profoundly impacted by AUD. The influence of the MD extends beyond its cortical connections, as it innervates and shares connections with brain regions essential for gustatory processing (Odegaard et al., 2025) and drug-seeking behavior (Vertes, 2002). The reciprocal projections with these brain regions enable the MD to orchestrate complex behaviors that could contribute to decisions surrounding alcohol misuse.

Although the MD has been relatively understudied in alcohol research, some evidence indicates that alcohol affects the MD in humans with AUD and preclinical models of alcohol exposure. Long-term damage to the MD and other midline thalamic regions has been documented in AUD individuals, such as those with Korsakoff’s syndrome (Halliday et al., 1994; Segobin et al., 2019). An analysis of tractography studies revealed overall degeneration of white matter tracts between the MD and PFC in AUD individuals (Segobin et al., 2019) that could underlie impaired behavioral flexibility and decision-making observed in AUD individuals. Cue-induced alcohol-seeking behavior has been linked to engagement of plasticity-related signaling mechanisms in the MD (Salling et al., 2017). Moreover, acute alcohol disrupts local field potential rhythms in the mouse MD (Choi et al., 2015), while alcohol dependence elevates MD serotonin transporter expression (Shibasaki et al., 2010) and increases c-Fos expression in PKC-ε KO knockout mice (Olive et al., 2001). While the MD is implicated in cognition and reward-based decision-making, its specific role in chronic alcohol-induced behavioral deficits, and whether it shows alcohol-induced neuroadaptations, remains relatively unexplored. Based on these findings, we hypothesized that chronic alcohol exposure in the mouse would lead to altered MD physiology, gene expression, and in vivo calcium signaling, and that signaling in the MD for alcohol drinking would be increased in CIE mice when given a choice for alcohol over other drinking solutions.

## 2. Experimental procedures

### 2.1. Animals

Adult male and female C57BL/6J mice (8-12 weeks old at time of experiment) were obtained from Jackson Laboratories (stock # 000664). Mice were single housed in standard cages with ad libitum access to food and water and were maintained on a 12-h reverse light cycle for at least 1 week prior to experiments. All experiments were conducted in accordance with the NIH Guide for the Care and Use of Laboratory Animals and the Institutional Animal Care and Use Committee at the Medical University of South Carolina.

### 2.2. Chronic intermittent ethanol (CIE) model of alcohol dependence

Ethanol vapor was delivered in Plexiglass inhalation chambers (Becker & Lopez, 2004; Padula et al., 2020). Ethanol levels in the chambers were monitored daily and adjusted to maintain 175-250 mg/dL blood ethanol concentrations. To confirm target levels were reached, retro-orbital blood was collected from a subset of mice each exposure day. Mice were placed into chambers for 16 h a day, followed by an 8-h withdrawal period. Each CIE cycle consisted of 4 days of ethanol exposure in the vapor chambers, followed by a 72-h withdrawal period. Immediately before being placed in chambers, CIE mice were injected with a loading dose of ethanol (1.6 g/kg) and pyrazole (1mmol/kg in 0.9% sterile saline; IP). Air control mice were handled in the same manner as CIE mice receiving pyrazole injections (1mmol/kg in 0.9% sterile saline; IP) immediately before being placed in chambers with room air.

### 2.3. Two-bottle choice preference test

Mice underwent 2-h two-bottle choice drinking sessions in their home cages five days/week (Becker & Lopez, 2004; Padula et al., 2015), beginning 3 h into the dark cycle. One bottle contained 15% (v/v) alcohol and the other contained H_2_O. During the alcohol-biased choice test (Lopez & Becker, 2025), the water bottle was replaced with a bottle of 3% (w/v) sucrose. The side of the alcohol-containing bottle was switched daily to control for habitual responses and side preference. Volume consumed was normalized to body weight. Preference for alcohol was calculated as the percentage of alcohol consumed over total fluid consumed. Mice were allowed to drink alcohol or H_2_O in their home cages for 2-6 weeks during baseline prior to CIE exposure then again for test drinking weeks following 1-2 weekly cycles of vapor exposure. The alcohol vs. sucrose test drinking session occurred after the 4^th^ cycle of CIE exposure.

### 2.4. c-Fos immunohistochemistry

Images of c-Fos staining in the MD were acquired from brain sections that were originally processed in a published study (Smith et al., 2020). In brief, mice underwent 5 alternating weeks of two-bottle choice (15% alcohol vs. H_2_O) home cage drinking (or no drinking) and CIE cycles (or Air control). Mice were perfused 2 h after withdrawal time points of interest: 0, 8, 24, 72 h, or 7 days. Brains were sliced into 40μm sections for c-Fos immunohistochemistry. Sections were permeabilized in 0.3% H_2_O_2_ for 15 min, followed by blocking in 2% normal donkey serum in PBS with 0.3% Triton X-100 for 1 h. Sections were incubated overnight at room temperature in primary antibody diluted in blocking solution (1:20,000 rabbit anti-c-Fos, EMD-Millipore, catalog # PC38). Sections were then incubated in secondary antibody (1:1000 biotinylated donkey anti-rabbit, Jackson Immuno Research) and then ABC (1:1000 Vector Elite Kit, Vector Labs) reagent for 45 min/each. C-Fos immunostaining was visualized by incubating sections in 0.025% 3,3’-diaminobenzidine (DAB), 0.05% nickel ammonium sulfate, and 0.015% H_2_O_2_. Images were captured at 4× magnification using an EVOS microscope. Two 40μm brain sections (approximately −1.34 and −1.70mm AP relative to Bregma) were acquired for each mouse. Two representative images capturing both the right and left hemispheres were acquired, and c-Fos-positive nuclei were counted on images in ImageJ using custom settings with a bandpass filter. Briefly, a region of interest (ROI) was created referencing a mouse brain atlas and was applied to each image to standardize image analysis across slices, and c-Fos+ cells were identified and counted automatically using ImageJ. c-Fos counts were normalized to the Air control group for each time point.

### 2.5. Gene expression analysis

MD tissue samples were collected at 8, 24, 72 h of withdrawal from 4 weekly cycles of CIE exposure. Adult male mice (n=7-8/group/time point) were deeply anesthetized via isoflurane, rapidly decapitated, and whole brains were quickly removed. One mm sections containing midline thalamus nuclei were obtained using a mouse brain matrix at 4°C, and the MD was collected from brain sections using a 1mm tissue punch and flash frozen immediately. Total RNA was purified using the ReliaPrep Kit (Promega, Cat # Z6210) following manufacturer’s instructions. Direct quantification of transcripts was performed using 100ng of total RNA on an nCounter MAX system (NanoString, Seattle, WA) using Mm NeuroPath V1.0 probes. RCC files were imported into nSolver 4.0 software and analyzed using the default criteria. Normalization was performed in nSolver using the geNorm algorithm with 9 housekeeping genes (*Ccdc127*, *Aars*, *Lars*, *Fam104a*, *Mto1*, *Csnk2a2*, *Supt7l*, *Tada2b*, and *Cnot10*). Functional annotation of the differentially expressed genes (DEGs) was performed in ToppGene (Chen et al., 2009) following our previously published methods (Mulholland et al., 2023; Womersley et al., 2024). To identify potential genes that are regulated by circadian rhythms, genes that differed across time in control mice were queried using the Circadian Genes in Eukaryotes database (Li et al., 2017).

### 2.6. Whole-cell slice electrophysiology

Brain slices containing the MD were prepared for whole-cell patch-clamp electrophysiology. Mice were briefly anesthetized with isoflurane and rapidly decapitated before quickly removing the brain. Brain slices (300µm) were collected using a Leica VT1000S vibratome containing cold oxygenated (95% O_2_, 5% CO_2_) sucrose cutting solution (200mM sucrose, 1.9mM KCl, 1.2mM NaH_2_PO_4_, 6mM MgCl_2_, 0.5mM CaCl_2_, 0.4mM ascorbate, 10mM glucose, and 25mM NaHCO_3_, adjusted to 305-315mOsm, pH 7.4). Slices were transferred into a holding chamber containing oxygenated artificial cerebrospinal fluid (aCSF; 125mM NaCl, 2.5mM KCl, 1.25mM NaH_2_PO_4_, 1.3mM MgCl_2_, 2.0mM CaCl_2_, 0.4mM ascorbate, 10mM glucose, and 25mM NaHCO_3_, adjusted to 290– 310mOsm, pH 7.4) at 34°C for 30 minutes. Slices were kept at room temperature for >30 minutes before recording. For recordings, a slice was placed in the recording chamber and perfused with 34°C aCSF at a flow rate of 2mL/min. Pipettes were prepared from thin-wall borosilicate glass electrodes and were pulled on a Sutter Instrument P97 Micropipette puller with tip resistances from 1.9 to 5.5MΩ. Current-clamp recordings were conducted in MD principle neurons using Axon MultiClamp 700B amplifiers (Molecular Devices) and Instrutech ITC-18 analog-digital converters (HEKA instruments) controlled by AxographX software. The patch pipette was filled with potassium gluconate internal solution (120mM KGluconate, 10mM KCl, 10mM HEPES, 2mM MgCl_2_, 1mM EGTA, 2mM NaATP, and 0.3mM NaGTP, adjusted to 294mOsm, pH 7.4). Firing was induced by applying a series of depolarizing and hyperpolarizing current steps (+/- 50, 70, 90, 110 pA) through the patch pipette. Events were filtered at 4kHz and digitized at a sampling rate of 10kHz. Recordings were made 2-7 h after euthanasia.

### 2.7. In vivo fiber photometry

To induce expression of a calcium biosensor, a viral vector construct for expression of GCaMP6f (pENN.AAV1.CamKII.GCaMP6f.WPRE.SV40; Addgene cat# 100834) was unilaterally injected in MD (200µL; 50 µL/min; AP: −1.58mm, ML: 0.44mm, DV: −3.2mm, relative to Bregma) followed by placement of a 2.5mm fiber optic ferrule containing a 400µm optical fiber (0.48NA) above the injection site coordinates (AP: −1.58mm, ML: 0.44mm, DV: −3.1mm). Mice were allowed to recover for at least 3 weeks prior to the start of experiments. During recording, GCaMP6f was excited using a collimated 470nm LED (Thorlabs), and signal from an equally powered 405nm LED was used for normalization to control for movement artifacts. Illumination power for each LED (5-30µW) was sinusoidally modulated at 531 Hz (405nm) and 211 Hz (470nm) and was passed through Doric fluorescent Mini Cubes. Emission light was focused onto a femtowatt photoreceiver (Newport, model 1672151; DC low setting) and sampled at 6.1kHz by a RZ5P lock-in digital processor controlled by photometry processing software (Synapse, Tucker-Davis Technologies, Alachua, FL, USA). Raw signals were pre-processed within Synapse software using a low-pass filter (3Hz) to remove system noise. Fiber optic patch cords were photobleached twice weekly overnight using a 405nm LED to reduce autofluorescence. The GCaMP6f signal was integrated in real-time with licking behavior using lickometer circuits (MedAssociates Inc., Fairfax, VT, USA) following our previous methods (Lebonville et al., 2025; Rinker et al., 2023).

Data were pre-processed using custom MATLAB functions (Mathworks). Because 405nm and 470nm signals show differences in photobleaching over time (Rinker et al., 2023), each channel was first fitted to a polynomial versus time and then subtracted from one another to calculate the Δ*F*/*F* time series. ‘Findpeaks’ function in MATLAB with a 2× threshold of the median average deviation of the normalized signal was used to identify GCaMP6f ‘events’.

### 2.8. Statistical analyses

Data are presented as mean ± SEM and were analyzed with GraphPad Prism (version 10.6.0; GraphPad, La Jolla, CA) or R software (v4.4.1) using unpaired t-tests or linear models with Tukey post-hoc, when appropriate. An α value of <0.05 was used to determine significance. For transcript analysis, DEGs were determined by linear regression analysis in nSolver software, and *p*-values were corrected using a 5% FDR with the Benjamini, Krieger, and Yekutieli approach in GraphPad Software (version 10.6.0). The GCaMP6f signal was extracted 8s before, during, and 8s after licking bouts, which were defined as lick frequency >4.5Hz, interlick interval <500ms, and duration >500ms based on the microstructure of rodent licking behavior (Boughter et al., 2007; Davis & Smith, 1992; Johnson et al., 2010). The GCaMP6f signal at the onset of licking bouts and the peak amplitude of the GCaMP6f signal within 3sec following bout termination were analyzed. Next, functional linear mixed models (FLMM) (Loewinger et al., 2024) were performed using R scripts to evaluate differences in the signals across this entire epoch. FLMM analysis allows for the comparison of the magnitude and timing of GCaMP6f signaling at each time point across treatment conditions while accounting for between-mouse differences (Loewinger et al., 2024). For FLMM analysis, signals were aligned to bout start, signals during bouts were interpolated (matching median bout duration using ‘interp1’ MATLAB function) so bout data were on equal time scales, and data were down-sampled to 10Hz for analyses. Data before/after bouts were not interpolated. While similar results were obtained when using mean bout duration for interpolation and interpolation has been previously applied to biosensor data (Amalyan et al., 2022; Markowitz et al., 2023), FLMM results during bouts should be interpreted with the understanding of data transformation to an equal time scale. Models that converged successfully and yielded lower AIC/BIC values were selected and included mouse-specific random functional intercepts and random functional slopes for solution [e.g., (Photometry_Signal ∼ Solution + (Solution|Mouse_ID); (Photometry_Signal ∼ Solution*Treatment + (Solution|Mouse_ID)]. Joint 95% confidence intervals (CIs) corrected for multiple comparisons for fixed effect β coefficients were considered statistically significant for the entire epoch when Y is different from 0.

## 3. Results

### 3.1. Alcohol dependence and c-Fos induction

To first assess how alcohol drinking and dependence affected MD c-Fos expression, we determined the number of c-Fos+ cells as a proxy for cellular activation using non-drinking and drinking mice exposed to CIE. A full description of methods and drinking data are described previously (Smith et al., 2020). Regardless of the drinking history, CIE exposure significantly reduced c-Fos+ cells at 2 h withdrawal in comparison with Air controls (treatment main effect: *F*_1, 19_=16.34, *p<*0.001; **Fig 1A,B**). In contrast, there was a significant increase in c-Fos+ cells at 10 h of withdrawal in CIE mice that returned to values below Air controls at 26 h (treatment main effect: *F*_1, 20_=7.12, *p*<0.05). No significant effects of CIE or drinking on c-Fos expression were observed at 74 h (main effect of treatment: *F*_1, 20_=0.643, *p*=0.43) or 7 d (treatment main effect: *F*_1, 19_=1.424, *p*=0.25) withdrawal. Notably, this pattern of MD c-Fos expression was similar to that reported for the lateral OFC, prelimbic cortex, and infralimbic cortex at 2 and 10 h withdrawal (Smith et al., 2020). However, the MD was the only region analzyed to show reduced c-Fos at the 26-h time point (treatment main effect: *F*_1, 19_=5.538, *p*<0.05), suggesting that CIE potentially alters the MD in a manner that does not merely mimic CIE-induced effects on its cortical connections.

**Fig. 1.**
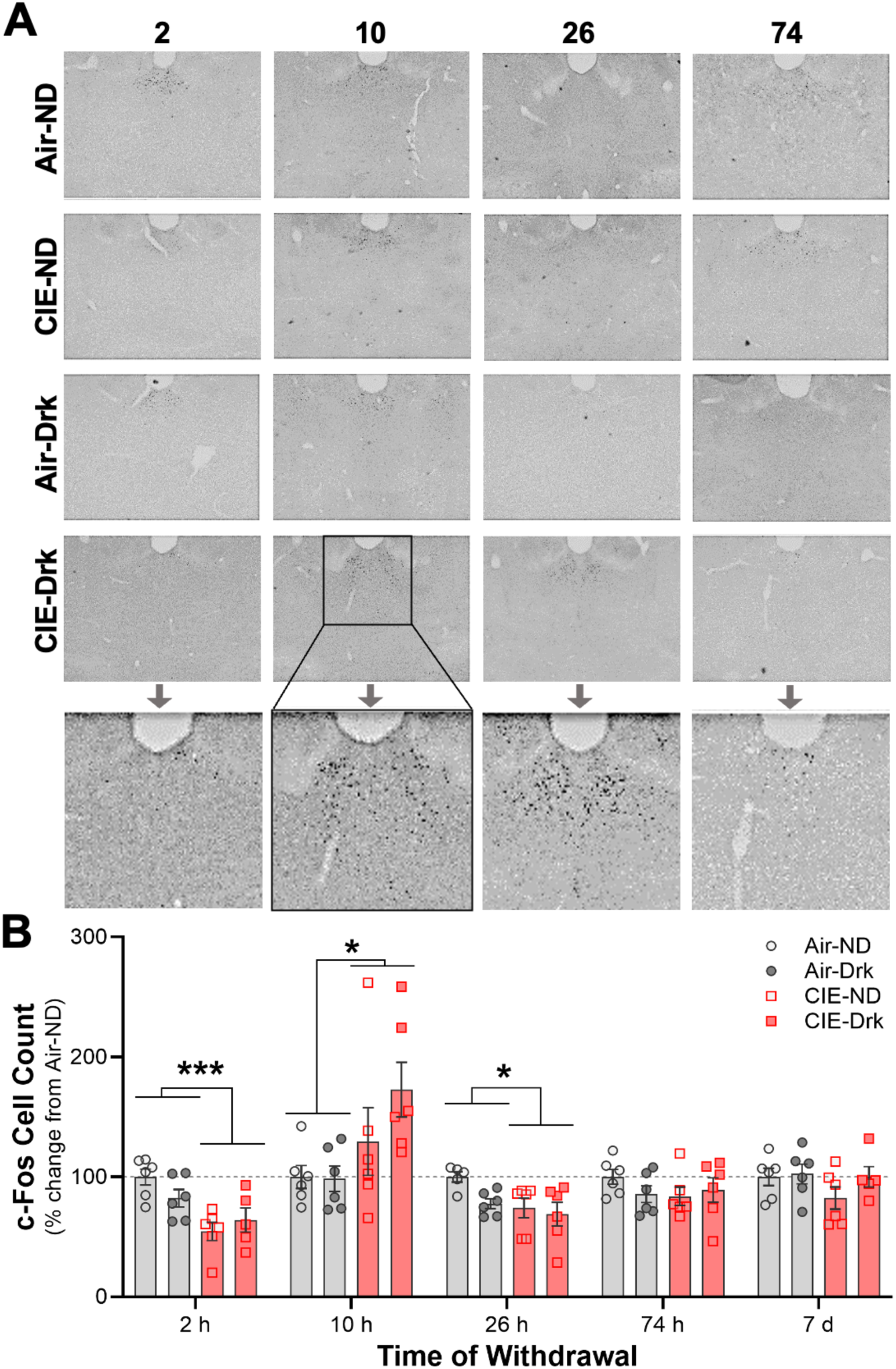
MD c-Fos expression in non-drinking (ND) and drinking (Drk) male mice that were exposed to chronic intermittent ethanol (CIE). (A) Representative images and (B) analysis of c-Fos+ cell count across time and treatment groups. *n*=5-6 mice/group/time point; \**p*<0.05, \*\*\**p*<0.001.

### 3.2. Differential gene expression

Because excessive drinking can lead to damage of the midline thalamic nuclei (Halliday et al., 1994), we determined whether CIE exposure affected gene expression using a panel that is enriched for neuropathology across five cell types: neurons, astrocytes, microglia, oligodendrocytes, and endothelial cells. Due to differences in tissue collection times, we first examined gene expression differences in Air controls to determine if there were genes regulated by circadian rhythms. There were 15 DEGs when comparing the 8-vs 24-h time points, and none when comparing 24-vs 72-h time points (**Fig 2A**). A search of the Circadian Genes in Eukaryotes database showed that 13 of these 15 DEGs have been linked to circadian rhythms in mice (**Table 1**).

**Fig. 2.**
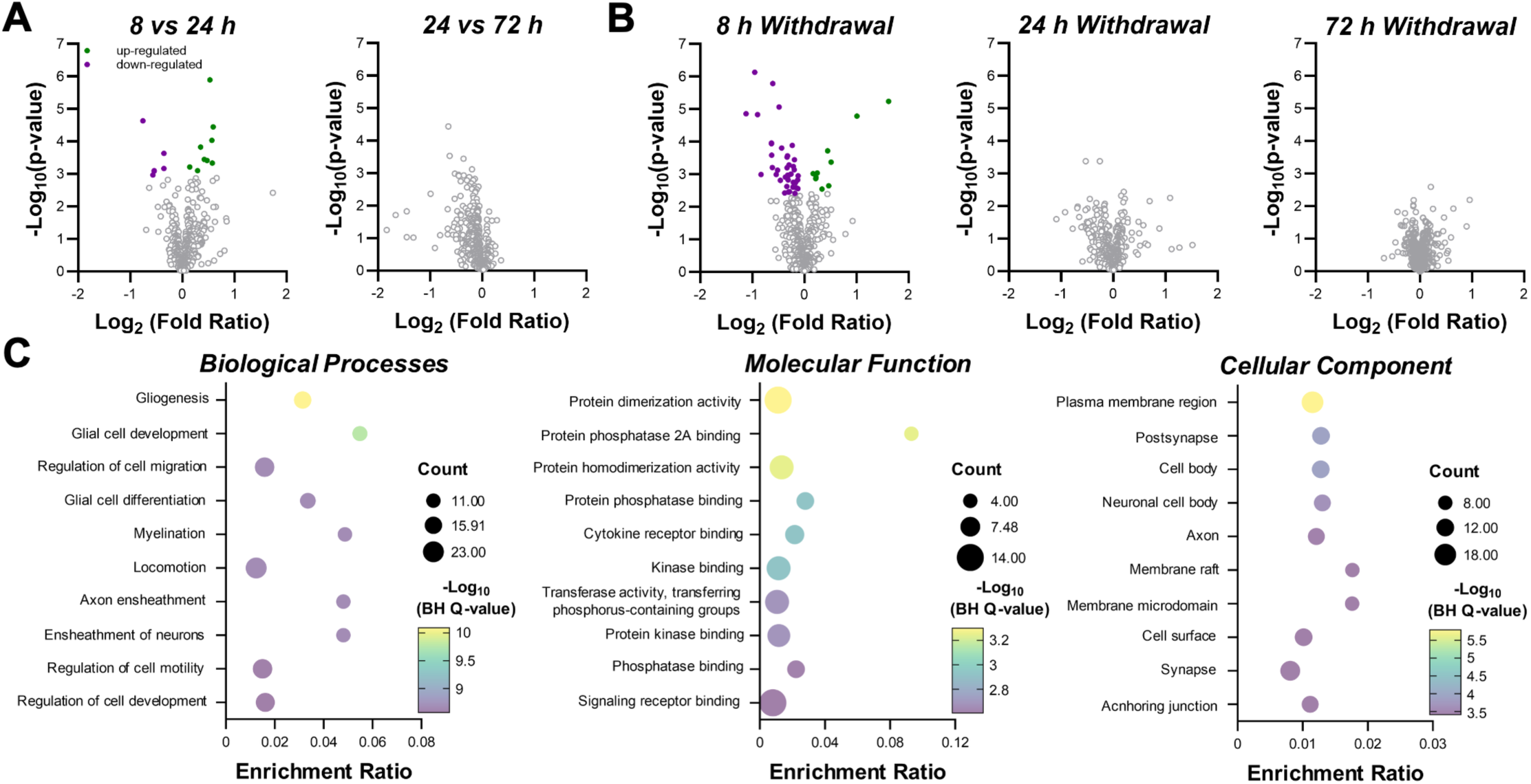
Differential gene expression analysis of neuropathology-related genes in CIE mice. Volcano plots comparing gene expression in (**A**) Air mice at different circadian time points and (**B**) Air vs CIE exposed mice at 3 times during withdrawal (*n*=7-8 mice/group/time). (**C**) Gene ontology enrichment analysis of MD differentially expressed genes of CIE mice.

**Table 1.**
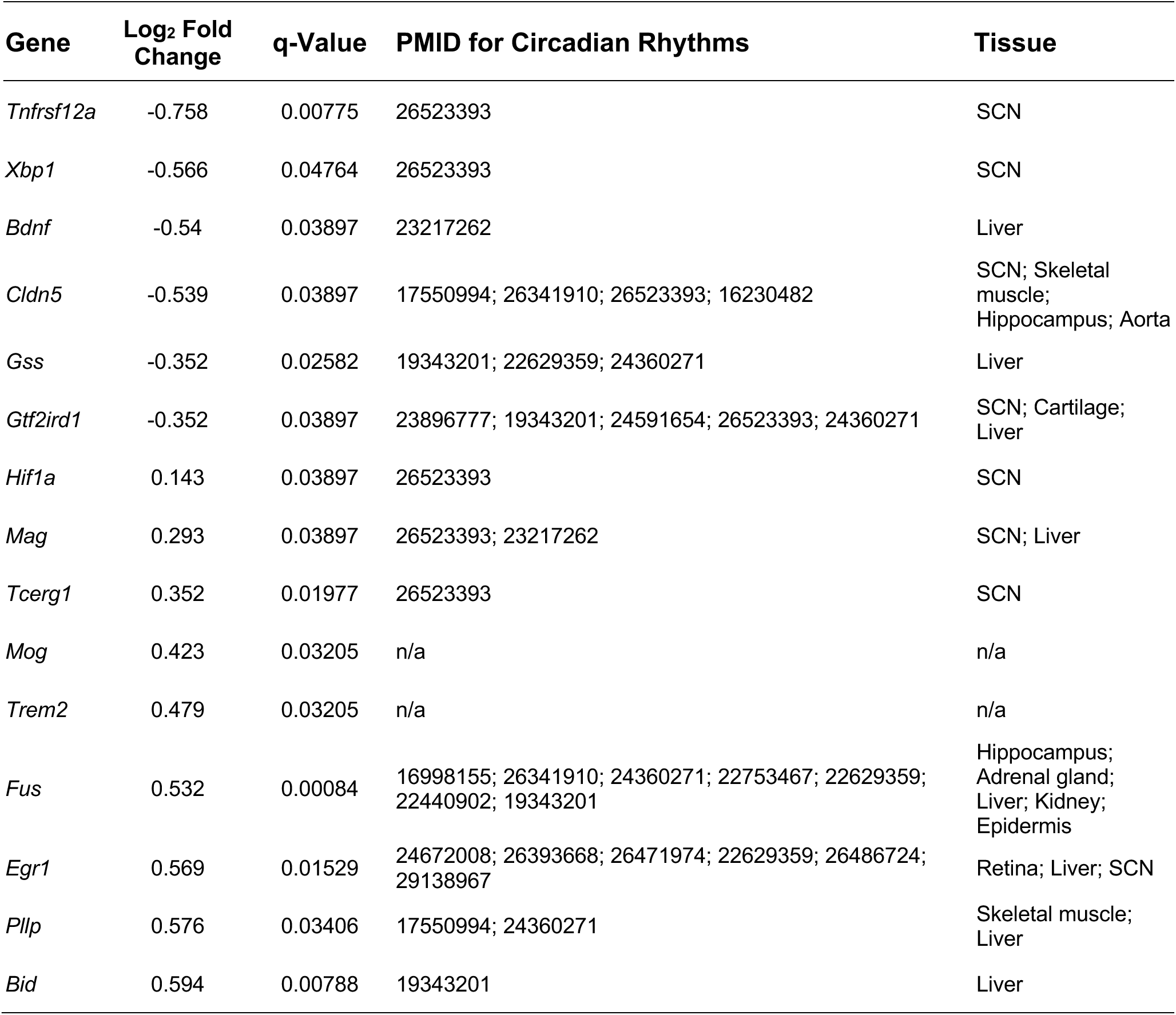
Time-dependent differentially expressed genes in the mediodorsal thalamus of control mice. SCN, Suprachiasmatic nucleus.

Differential expression analysis revealed that CIE dysregulated 49 MD genes at 8 h into withdrawal (**Table 2**; **Fig 2B**), the majority of which (39) were down-regulated. In comparison, CIE exposure did not significantly alter genes at 24 or 72 h of withdrawal. Gene Ontology enrichment analysis of the DEGs at 8 h of withdrawal showed that CIE disrupted processes related to glial function, axonal myelination, protein dimerization, and binding for protein phosphatase, kinase, cytokine receptors, and signaling receptors (**Fig 2C**). DEGs localized to different subcellular compartments, including synapses and axons, suggesting possible physiological adaptations in CIE mice.

**Table 2.**
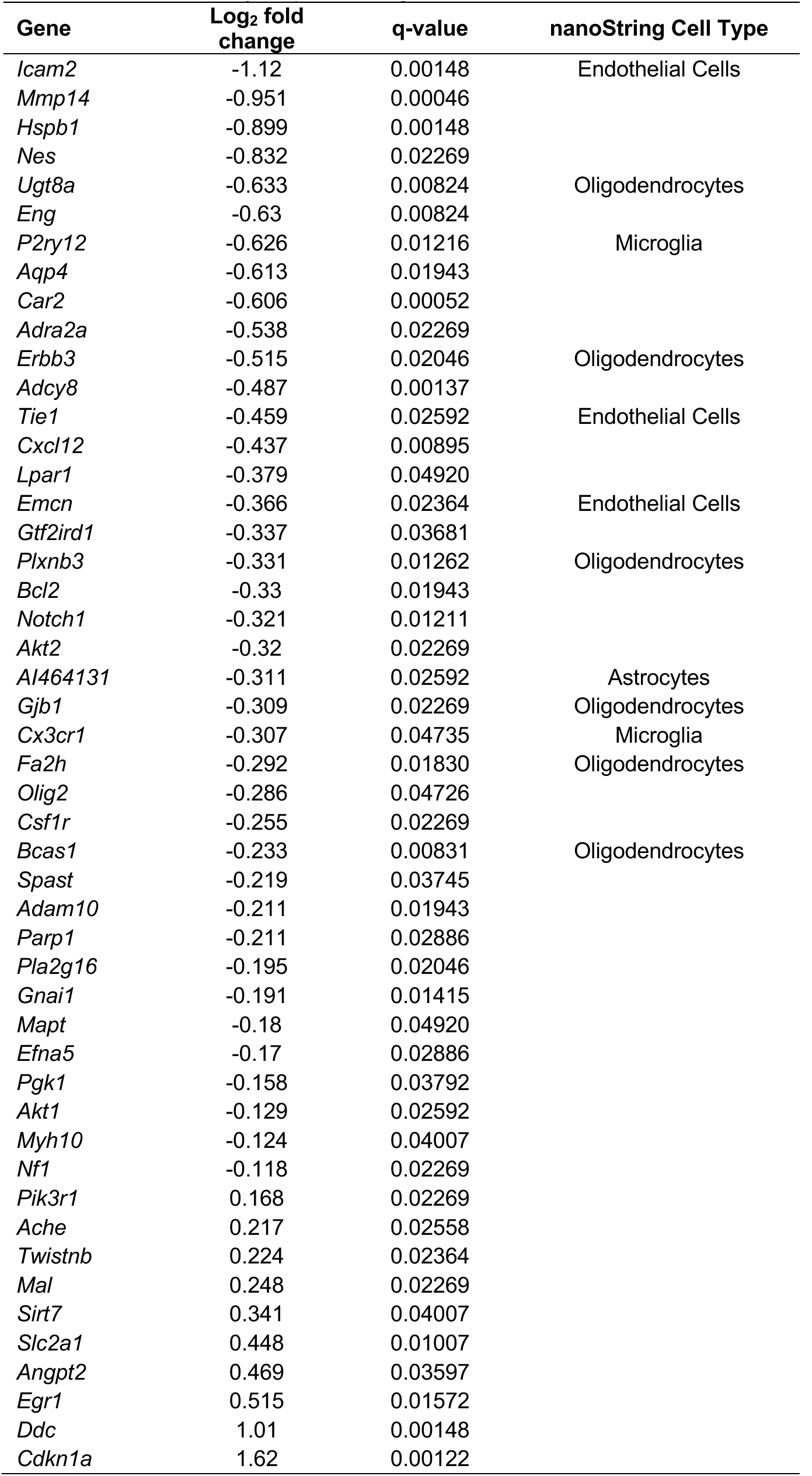
Differentially expressed genes in the mediodorsal thalamus of CIE exposed mice.

### 3.3. Physiological adaptations

While we identified MD adaptations during early withdrawal from CIE exposure, it is unclear if there are changes in MD physiology in CIE mice. To address this question, we performed slice electrophysiology recordings in MD neurons at 72 h of withdrawal. This time was selected because mice show increased drinking when given alcohol access in their home cages (see **Fig 4A** and **Fig 5B**). Evoked action potentials were elicited during hyperpolarizing and depolarizing current injections. As shown in **Fig 3A,B**, CIE exposure significantly increased evoked firing of MD neurons in comparison with Air mice regardless of the polarity of the current injection (treatment main effect: *F*_1,200_=43.25, *p*<0.0001; *n*=7-8 mice/group; 3-6 cells/mouse). Analysis of the biophysical membrane properties revealed a significant decrease in the threshold for action potential firing (*t*_61_=2.049, *p*<0.05; **Fig 3C**) and a trend for a reduction in the medium after-hyperpolarization amplitude (*t*_61_=1.747, *p*=0.08) in CIE exposed mice. Other measures of membrane properties were not affected by CIE (*p*>0.1; **Fig 3C-J**). Thus, CIE exposure increased excitability of MD neurons.

**Fig. 3.**
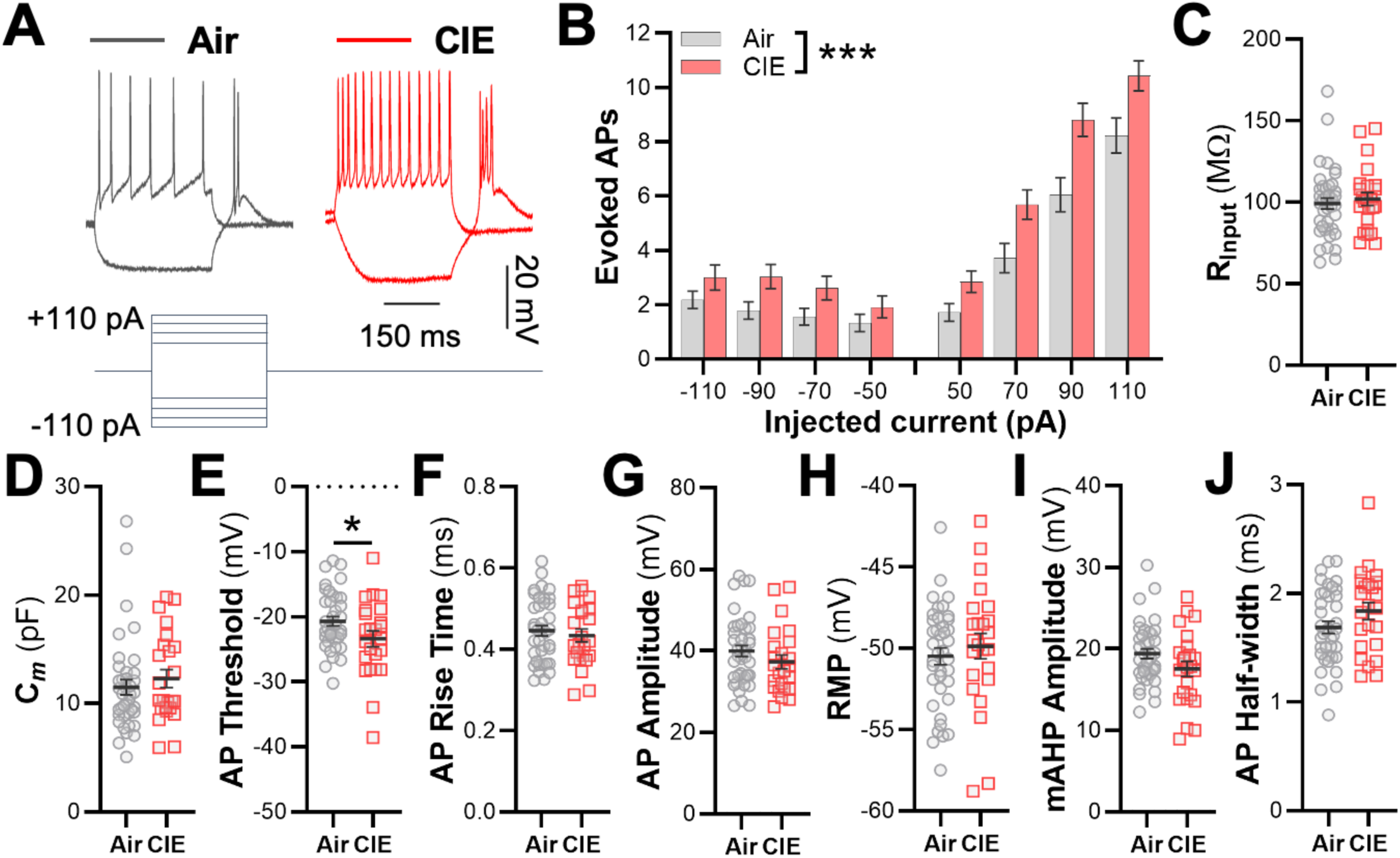
Evoked action potentials (AP) recorded from MD neurons in control and CIE male mice. **A)** Representative traces of hyperpolarizing and depolarizing current steps. **B)** Evoked firing of MD neurons at 72 h withdrawal from CIE exposure. **C-J)** Biophysical membrane properties of MD neurons. *n*=7-8 mice/group; 3-6 cells/mouse; \**p*<0.05, \*\*\**p*<0.001.

**Fig. 4.**
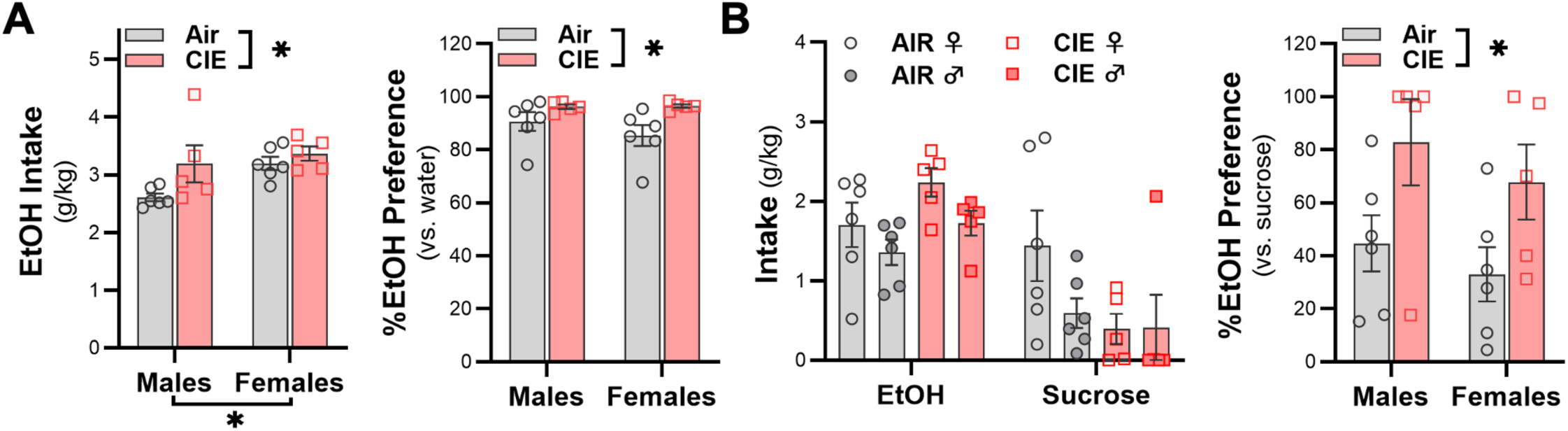
Drinking in CIE mice when given a choice between alcohol and water or sucrose. **A)** Alcohol drinking and preference over water in CIE exposed male and female mice. **B)** Alcohol drinking and preference over sucrose. *n*=5-6 mice/sex/group; \**p*<0.05.

**Fig. 5.**
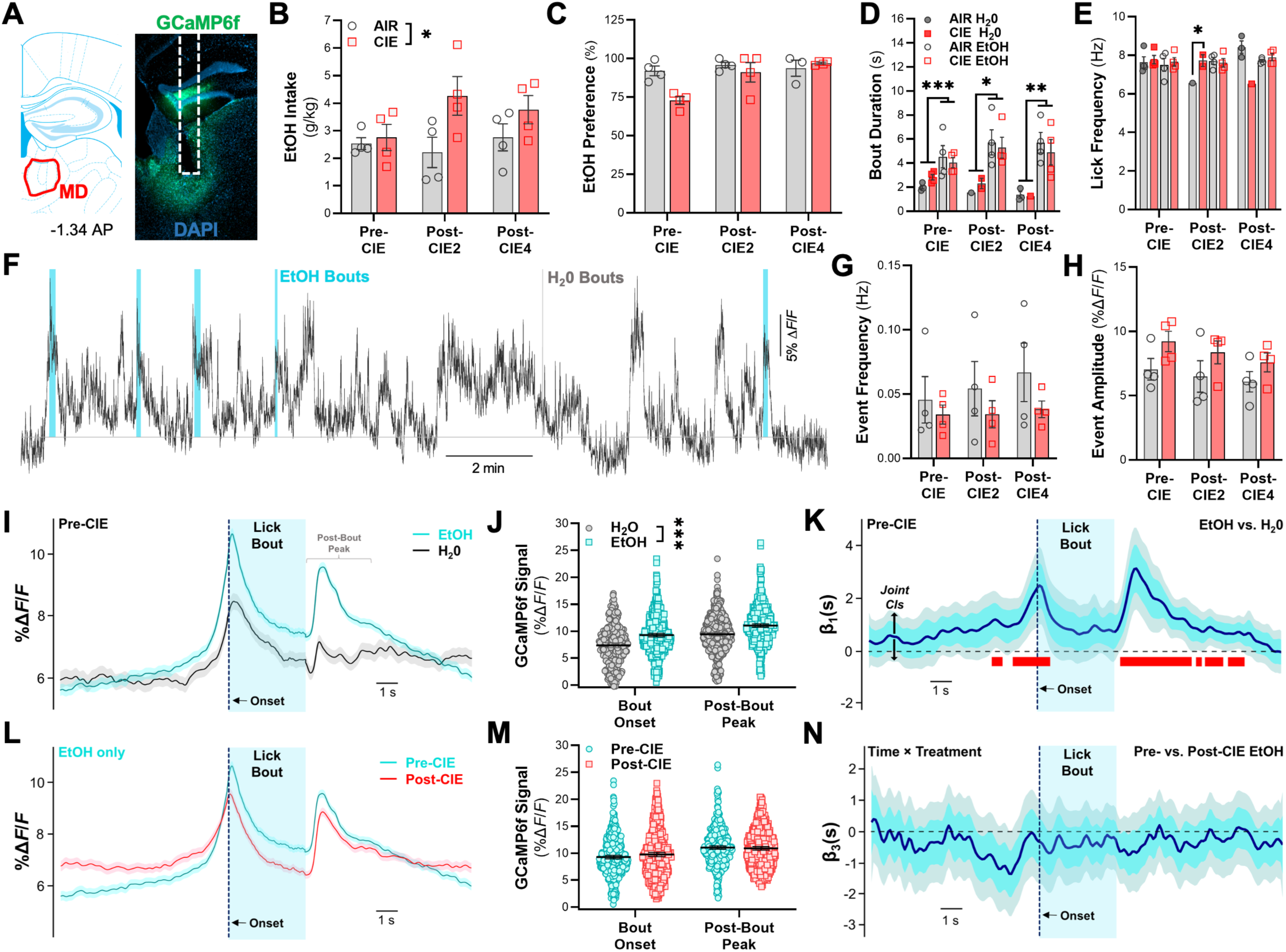
Alcohol consumption and GCaMP6f signaling in CIE male mice. **A)** Optical ferrule and GCaMP6f expression in the MD for fiber photometry recordings. **B)** Alcohol intake and **C)** preference in CIE-exposed mice versus Air controls across three time points. **D)** Bout duration and **E)** lick frequency during choice tests for alcohol and water. **F)** Representative GCaMP6f signal in the MD during alcohol and water drinking. Analysis of GCaMP6f event **G)** frequency and **H)** amplitude. **I)** Averaged photometry traces, **J)** GCaMP6f signal surrounding licking bout, and **K)** FLMM analysis depicting functional β coefficient estimate plots when comparing alcohol vs. water. **L)** Averaged photometry traces, **M)** GCaMP6f signal surrounding licking bout, and **N)** FLMM analysis depicting functional β coefficient estimate plots for alcohol drinking when comparing pre- and post-CIE exposure. Red bars denote significance when joint corrected confidence intervals (CIs, shown in black arrows) do not contain a zero value. *n*=4 mice/group; \**p*<0.05.

### 3.4. Alcohol-biased choice behaviors

Little research has focused on how alcohol misuse alters neural activity to modulate alcohol-biased choice behaviors in mice. To address this, we first replicated a previous study demonstrating alcohol-biased choice in the CIE mouse model of dependence (Lopez & Becker, 2025). Male and female mice were allowed to drink alcohol or water in the two-bottle choice model during a 2-week baseline period prior to exposure to Air or CIE. Following CIE exposure, mice were given access to sipper tubes containing alcohol and either water or sucrose. As expected, CIE exposure increased alcohol intake (treatment main effect: *F*_1,18_=4.77, *p<*0.05; *n*=5-6 mice/sex/group) and preference (treatment main effect: *F*_1,18_=7.86, *p*<0.05) (**Fig 4A**). Alcohol drinking was also higher in female compared with male mice (sex main effect: *F*_1,18_=4.92, *p*<0.05). When given the choice between alcohol and sucrose, preference for alcohol reduced from >85% to <45% in Air controls, whereas preference for alcohol remained relatively high (65-80%) in CIE mice (**Fig 4B**). Analysis revealed a main effect of treatment (*F*_1,18_=8.23, *p*<0.05) where preference for alcohol when sucrose was available as an alternative was higher in CIE mice.

### 3.5. Elevated MD signaling during alcohol drinking

In a separate cohort of mice, calcium signals and fiber photometry were used to investigate the impact of CIE on MD activity during drinking (**Fig 5A**). Male mice (*n*=4/group) were given bottles containing 15% alcohol and water across multiple time points: before CIE exposure (Pre-CIE) and after two (Post-CIE2) or four (Post-CIE4) back-to-back CIE cycles. Ethanol intake increased in CIE-exposed (treatment main effect: *F*_1,9_=9.49, *p*<0.05; **Fig 5B**). While there was a significant main effect of time for preference (*F*_2,50_=4.37, *p*<0.05; **Fig 5C**), post-hoc comparisons of different time points were not significant.

Drinking bout duration differed depending on the time and solution offered (time×solution interaction: *F*_2,1087_=3.43, *p*<0.05; **Fig 5D**). Specifically, post-hoc analyses showed that bout duration for alcohol was longer than bouts for water before CIE exposure (*p*<0.0001) and after 2 (*p*<0.05) and 4 weeks of CIE exposure (*p*<0.01). Duration of alcohol drinking bouts increased after Post-CIE2 (*p*<0.01) and Post-CIE4 (*p*<0.05) test sessions compared with baseline. There was a significant interaction for lick frequency (group×time×solution interaction: *F*_1,1089_=4.69, *p*<0.01; **Fig 5E**), and post-hoc tests revealed a difference between water and alcohol during the Post-CIE2 test (*p*<0.05). A representative photometry recording during an alcohol vs. water drinking session is shown in **Fig 5F**. For GCaMP6f event frequency, analysis revealed a significant main effect of time (*F*_2,51_=4.44, *p*<0.05; **Fig 5G**), but post-hoc analyses did not show significant differences. Analysis of GCaMP6f event amplitude did not reveal significant changes across treatment or time (**Fig 5H**). As shown in **Figs 5F** and **6F**, licking occurred when the signal was elevated above baseline. Averaged GCaMP6f traces surrounding licking bouts are shown in **Fig 5I,L**. The GCaMP6f signal at the onset and peak following licking bouts was significantly higher for alcohol compared to water during Pre-CIE drinking sessions (solution main effect: *F*_1,1686.7_=227.99, *p*<0.001; **Fig 5J**). FLMM analyses (akin to a paired t-test comparing water and alcohol bouts at each timepoint with multiple comparison corrections) showed the signal was significantly higher on alcohol bouts than water bouts around the start and following drinking bouts. This suggests alcohol-specific calcium activity elevations in the MD (**Fig 5K**). The signal surrounding alcohol bouts remained stable following CIE exposure and did not differ between Air and CIE groups (treatment main effect: *F*_1,7.83_=0.09, *p*>0.7; **Figs 5M**; FLMM time×treatment interaction: Joint CIs contain 0; **Fig 5N**). Because mice consumed little to no water post-CIE, water drinking bouts were excluded from analysis.

**Fig. 6.**
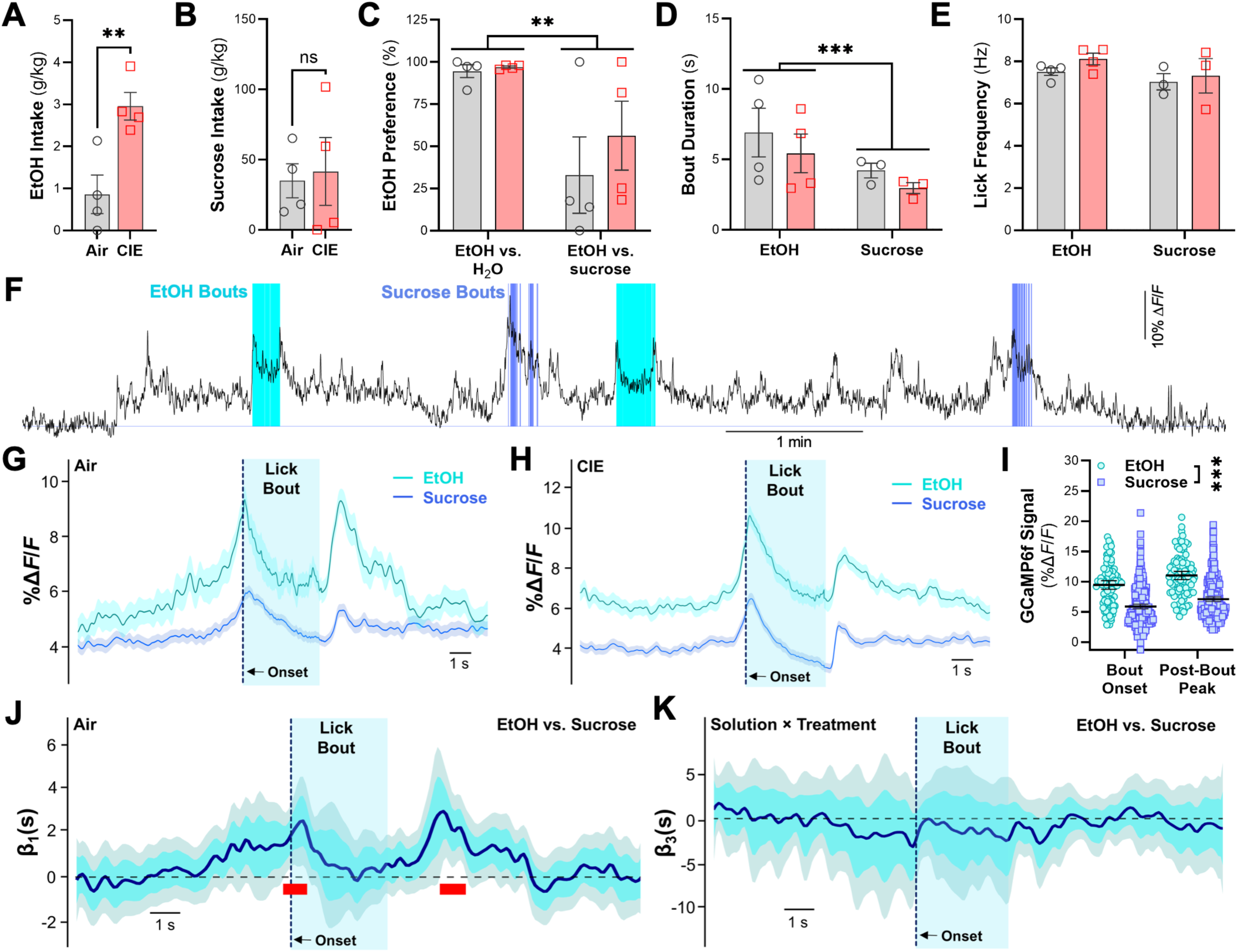
Drinking behavior and MD activity in CIE mice when given a choice between alcohol and sucrose. **A)** Alcohol and **B)** 3% sucrose intake during the choice test. **C)** Alcohol preference in the presence of a choice for alcohol and water or alcohol and sucrose. **D)** Bout duration and **E)** lick frequency during alcohol vs. sucrose choice sessions. **F)** Example fiber photometry recording of MD GCaMP6f activity during alcohol vs. sucrose choice drinking. **G,H)** Averaged GCaMP6f traces surrounding licking bouts in Air and CIE mice. **F)** Functional coefficient estimates from FLMM analysis of MD signaling for **I)** alcohol vs. sucrose and **J)** solution×treatment interaction. Red ticks denote significance when joint corrected CIs do not contain zero. *n* = 4 mice/group; \**p*<0.05, \*\**p*<0.005.

After completing alcohol vs. water drinking sessions post-CIE4 exposure, mice underwent a two-bottle choice test for alcohol vs. 3% sucrose. Alcohol intake in CIE mice was significantly higher than Air controls in sucrose choice session (*t*_6_=3.711, *p*<0.01; **Fig 6A**), whereas sucrose intake was similar between groups (*t*_6_=0.247, *p>*0.8; **Fig 6B**). Alcohol preference significantly decreased in the alcohol vs. sucrose choice session compared to when water was the alcohol alternative (session main effect: *F*_1,9_=18.96, *p*<0.05; **Fig 6C**). Analysis of bout duration and lick frequency indicated a significant main effect of solution (duration: *F*_2,623_=7.741, *p*<0.001; frequency: *F*_2,623_=5.072, *p*<0.001; **Fig 6D,E**), where alcohol was greater than sucrose for both measures. A representative photometry recording during an alcohol vs. sucrose choice session is shown in **Fig 6F**, and representative GCaMP6f traces surrounding licking bouts for alcohol and sucrose in Air and CIE mice are shown in **Fig 6G,H**. When comparing the signal for alcohol and sucrose during choice drinking, the calcium signal was briefly but significantly increased at the start of and after alcohol bouts (solution main effect: *F*_1,775.09_=81.75, *p*<0.001; **Fig 6I**; FLMM results in **Fig 6J**). The FLMM interaction between solution and treatment did not reveal differences in GCaMP6f surrounding drinking (**Fig 6K**). Together, photometry studies revealed no effect of CIE on MD signaling, although there are differences in the photometry signal within-animal between solutions with an elevated signal for alcohol compared to alternative solutions.

## 4. Discussion

We explored neuroadaptations and in-vivo signaling in the MD of CIE mice because of MD involvement in reward-guided and cognitive behaviors and its interconnections with cortical brain and gustatory regions. We found time-dependent differences in c-Fos expression, physiology, and gene expression during acute (<24 h) and protracted (3-7 days) withdrawal from chronic alcohol exposure in mice. Functional changes in membrane excitability of MD neurons occurred at a time point where CIE mice consumed more alcohol than control mice. In this study, CIE male and female mice also showed a strong preference for alcohol over a natural reward when given a choice between the two solutions in a home cage drinking model. Using calcium signaling and fiber photometry during home cage drinking sessions, we observed increased activity of MD neurons surrounding drinking bouts, and the signal for alcohol was higher than other solutions. Overall, this study shows physiological and genetic adaptations in the MD using a dependence model that increases alcohol drinking and shifts preference toward alcohol over a naturally rewarding solution.

To map activity changes in the MD across acute and protracted withdrawal from CIE exposure, we first measured c-Fos expression in control and CIE mice at five different time points. Similar to its cortical partners, c-Fos was reduced after removal from vapor chambers while mice were still intoxicated and increased at the peak of withdrawal hyperexcitability. In contrast, early withdrawal from an alcohol liquid diet did not activate c-Fos in the MD (Olive et al., 2001) in wildtype mice perhaps due to the difference in the severity of alcohol exposure produced by a liquid diet versus the vapor model. Although c-Fos was not reduced in withdrawn wild-type mice in that study, it was increased in PKC-ε KO mice. This suggests a possible genetic component to MD activation during acute alcohol withdrawal, although *Prkce* expression was not altered in the current study, in contrast to reports of increased expression in other model systems and brain regions (Hanim et al., 2023; Kumar et al., 2016; McClintick et al., 2014; Messing et al., 1991). When compared to its interconnected cortical regions and 24 other brain regions analyzed in Smith et al. (Smith et al., 2020) that showed no change or elevated c-Fos at 24h of withdrawal, we observed a reduction in c-Fos expression in the MD, demonstrating a unique response of the MD during withdrawal from CIE exposure in C57BL/6J mice. c-Fos expression in the MD returned to control values at 3-7 days of withdrawal, and this was also unique compared to the reduced expression observed in the prelimbic and infralimbic cortices at these later withdrawal time points (Smith et al., 2020). Thus, these data show that the MD is differentially activated across acute and more prolonged alcohol withdrawal, and these changes do not always parallel the response pattern in its interconnected cortical regions.

Next, we interrogated physiological adaptations in MD neuron firing in CIE mice. While c-Fos expression did not differ between control and CIE exposed mice at 3 days of withdrawal, excitability was increased in CIE mice following both depolarizing and hyperpolarizing current steps. Apart from action potential threshold, which decreased in CIE mice, other biophysical membrane properties of MD neurons were not altered. Whereas evoked action potentials during depolarizing pulse are driven by Na^+^ spikes, the rebound burst firing following hyperpolarization pulses in thalamic neurons are mediated by low-voltage activated T-type Ca^2+^ channels. In mouse nucleus reuniens of thalamus, CIE exposure increased low-threshold bursts and T-type Ca^2+^ channel gene expression and shifted T-type Ca^2+^ channel gating properties during acute (<1 d), but not protracted (>1 d) withdrawal (Graef et al., 2011). Compared with the reuniens, physiological adaptations in rebound burst firing of MD neurons persisted for a longer duration during withdrawal, suggesting unique adaptations in the MD compared with other midline thalamic nuclei. The mechanisms underlying the increased excitability of Na^+^ and Ca^2+^ spikes in MD neurons from CIE mice are unknown at this point, but may be driven by multiple disrupted processes, including shifts in T-type Ca^2+^ channel expression or gating properties that will be explored in future studies.

Examination of a neuropathology gene panel revealed differential expression of genes in CIE mice at 8 h of withdrawal, but not at later time points. Enrichment analysis clustered the DEGs into processes related to glial function, axonal myelination, and protein binding. These results add to a growing literature showing that alcohol drinking and dependence induce genetic and morphological changes in microglial (Marshall et al., 2020; Marsland et al., 2023; Melbourne et al., 2024; Nelson et al., 2021; Peng et al., 2017; Salem et al., 2024; Schrank et al., 2024; Warden et al., 2020) and astrocytes (Bull et al., 2014; Erickson et al., 2018; Kastner-Blasczyk et al., 2025; Nentwig et al., 2024). None of the genes included in the panel as putative ‘neuronal’ markers were differentially expressed. Further supporting predominantly glial adaptations, neither ‘endothelial cell function’ nor ‘apoptosis/cell death’ were identified as top terms in the enrichment analysis. Thus, these results from the neuropathology panel suggest that the CIE model does not produce neural damage or endothelial cell dysfunction in the MD as measured by gene expression. Despite these preclinical findings, there is profound neuronal loss in MD in AUD individuals with Wernicke-Korsakoff syndrome (Halliday et al., 1994), and a recent tractography study using diffusion tensor imaging and high-resolution MRI identified degeneration of white matter tracts interconnecting the MD and the PFC in individuals with AUD (Segobin et al., 2019). It is also possible that more subtle indicators of neuronal degeneration went undetected in our study. However, we did observe dysregulation of cell firing that might reflect changes in genes that control axonal processes and myelination. It is also possible that CIE exposure produces unique genetic adaptations in specific circuits (e.g., MD→PFC) that were not identified using our bulk tissue approach. Thus, while these studies identified adaptations in transcripts for specific biological processes, further studies are necessary to examine the full extent of genetic and cell-type specific changes that occur across time in the MD of CIE mice.

In addition to the DEGs identified in CIE mice, we also identified DEGs when comparing different time points in the circadian rhythm of alcohol-naïve mice. Searching the Circadian Genes in Eukaryotes database confirmed that the majority (13 of 15) of these DEGs have been previously linked to circadian rhythms in mice, partially validating results from the gene panel analysis using bioinformatics sources. Many of the genes were identified as having a role in circadian rhythms based on studies in the liver and superchiasmatic nucleus, an organ and brain region with known pacemaker actions for controlling metabolic and physiological processes. Evidence suggests that the MD is involved in processes related to several aspects of circadian rhythms, including arousal (Sriji et al., 2021), sleep (Choi et al., 2015; Schreiner et al., 2022; Shin et al., 2023), and time perception (Lusk et al., 2020), although the exact role of the MD DEGs in controlling physiological processes important for circadian rhythms is unknown. Interestingly, expression of two of the circadian-related DEGs (*Egr1* and *Gtf2ird1*) were also differentially expressed in the MD of CIE mice. Individuals with AUD often shown signs of circadian misalignment, including disrupted sleep cycles, body temperature, and altered molecular rhythms and neuroendocrinological functions (Davis et al., 2018; Koob & Colrain, 2020; Meyrel et al., 2020; Perreau-Lenz & Spanagel, 2015; Spanagel et al., 2005). Our findings suggest that these DEGs in the MD could control some aspects of circadian rhythms, and that the transcription factors *Egr1* and *Gtf2ird1* are potential candidate genes for perturbation of circadian alignment in AUD.

In a home cage choice model for alcohol or a non-drug reward (Lopez & Becker, 2025), we showed that preference for alcohol remained high in male and female C57BL/6J mice following CIE exposure. In both sexes, alcohol intake and preference increased in CIE mice and preference for alcohol over water in both groups and sexes was >80%. However, when given a choice between alcohol and sucrose in their home cage, preference for alcohol in the control mice dropped to below 45%, demonstrating that the CIE dependence model induces a high preference phenotype for alcohol over a non-drug reward. Consistent with the preference for alcohol over social rewards in male and female rats (Augier et al., 2023; Marchant et al., 2023), we showed that induction of alcohol dependence maintains preference for alcohol over sucrose in both sexes providing another preclinical model to assess alcohol-biased choice behaviors that have been described in AUD (Hogarth & Field, 2020). Using fiber photometry recordings of calcium signaling in the MD during home cage drinking, we showed MD activity increases around bouts for alcohol, water, and sucrose. MD signaling for alcohol bouts was significantly higher than bouts for water, and signal for alcohol remained elevated after CIE exposure as there was a lack of significant differences between-group in the FLMM analysis of the photometry signal. In control and CIE mice given a choice between alcohol and sucrose, there were small but significant increases in the MD signal during drinking bouts for alcohol versus sucrose. These results are consistent with the idea that the MD encodes taste identify (Odegaard et al., 2025) and cost-benefit decision-making (Ding et al., 2024). Thus, in a voluntary drinking model using different solutions, these studies demonstrated a unique within-mouse signal in the MD during alcohol consumption and no significant differences in signal response to alcohol between control and CIE mice suggesting an acute association between alcohol consumption and the photometry signal.

In summary, alcohol dependence in mice produces physiological and transcriptomic adaptations in the MD that coincide with excessive drinking and biased choice behaviors for alcohol over a natural reward. We also observed a unique signature of MD calcium activity for alcohol drinking that was maintained in CIE mice given choices between alcohol and water or a naturally rewarding solution.

### Author Disclosure and Role of Funding

Supported in part by the Charleston Alcohol Research Center (P50 AA010761), MUSC Post-Baccalaureate Research Education Program (R25 GM113278), the Summer Undergraduate Research Program in Neuroscience of Addiction (R25 DA033680), Training in Alcohol Research Program (T32 AA007474), the Translational Science Laboratory Core, Hollings Cancer Center Shared Resources, Medical University of South Carolina (P30 CA138313), and NIH grants K01 AA025110 (JAR) and R01 AA023288 (PJM).The NIH had no further role in study design; in the collection, analysis and interpretation of data; in the writing of the report; and in the decision to submit the paper for publication.

## Contributions

JCM completed analyses and wrote the first draft of the manuscript. KCR, KAC, MD, AS, EM, CLL, SNR, and PJM, collected data and completed analyses. MH and GL provided code and analytical guidance. KCR, JJW, RJS, MFL, HCB, PJM, and JAR designed the studies. All authors contributed to and have approved the final manuscript. All authors declare that they have no conflicts of interest.

## Notes

### Competing Interest Statement

The authors have declared no competing interest.

### Summary of Updates

Figure 4 was revised and new analysis included. Figure 5 and 6 have revised FLMM analyses. Authors were updated to include additional authors based on updated analyses and contributions.

